# An Omentum-inspired 3D PEG Hydrogel for Identifying ECM-drivers of Drug Resistant Ovarian Cancer

**DOI:** 10.1101/560482

**Authors:** Elizabeth A. Brooks, Maria F. Gencoglu, Daniel C. Corbett, Kelly R. Stevens, Shelly R. Peyton

## Abstract

Ovarian cancer (OvCa) is a challenging disease to treat due to poor screening techniques and late diagnosis. There is an urgent need for additional therapy options, as patients recur in 70% of cases. The limited availability of clinical treatment options could be a result of poor predictions in early stage drug screens on standard tissue culture polystyrene (TCPS). TCPS does not capture the mechanical and biochemical cues that cells experience *in vivo*, which can impact how cells will respond to a drug. Therefore, an *in vitro* model that captures some of the microenvironment features that the cells experience *in vivo* could provide better insights into drug response. In this study, we formed 3D multicellular tumor spheroids (MCTS) in microwells, and encapsulated them in 3D omentum-inspired hydrogels. SKOV-3 MCTS were resistant to Paclitaxel in our 3D hydrogels compared to a monolayer on TCPS. Toward clinical application, we tested cells from patients (ovarian carcinoma ascites spheroids (OCAS)), and drug responses predicted by using the 3D omentum-inspired hydrogels correlated with the reported pathology reports of those same patients. Additionally, we observed the presence of collagen production around the encapsulated SKOV-3 MCTS, but not on TCPS. Our results demonstrated that our 3D omentum-inspired hydrogel is an improved *in vitro* drug testing platform to study OvCa drug response for patient-derived cells, and helped us identify collagen 3 as a potential driver of Paclitaxel resistance.

## Introduction

Ovarian cancer (OvCa) is the fifth deadliest cancer for women, and the deadliest gynecological disease for women overall, resulting in an estimated 14,070 deaths in the United States in 2018.^1^ The high mortality rate of OvCa is due to late detection, inadequate screening techniques, and a lack of effective second line therapies.^2^ OvCa is typically detected very late, partially because the disease is asymptomatic until Stage III,^3^ where the cancer cells are no longer confined to the ovaries. At the time of staging laparectomy, metastases that have spread through the peritoneum are present in 70% of patients.^3^ Tumor cells spread into the peritoneal fluid, where they form ovarian carcinoma ascites spheroids (OCAS), and can attach onto the abdominal peritoneum or omentum.^4, 5^ Prognosis for OvCa patients has improved little in the last three decades since the introduction of platinum-based treatments.^6^ Although first line therapy results in complete remission in 40-60% of patients, over 70% of those patients relapse within two years, and there are no effective second line treatments.^7, 8^ Additionally, to our knowledge, only four targeted drugs have been approved by the Food and Drug Administration (FDA) in the last ten years for OvCa. There is a clear need for innovation in drug discovery to develop additional OvCa treatments to improve patient outcomes.

A possible reason for the inadequate success in OvCa drug discovery is that current pre-clinical *in vitro* screening methods are not good predictors of clinical success of drug candidates. Many hypothesize that this is largely because *in vitro* screening uses cell lines grown on tissue culture polystyrene (TCPS).^9, 10^ The TCPS surface has no resemblance to the *in vivo* microenvironment chemically, physically, or topologically.^11^ Cancer cells grown in this environment have different morphologies, phenotypes, cellular signaling, and drug responses than cells found *in vivo*.^12^ Three-dimensional (3D) culture models are increasingly used in basic science applications because they offer environments that better resemble *in vivo* conditions,^13, 14^ and they can be tuned to model different tissues.^15, 16^ Most 3D models consist of hydrogels or similar biomaterials that have a 3D structure inside which cancer cells can be grown.^17–20^ These materials can be functionalized with peptides to mimic cell-extracellular matrix (ECM) interactions, ECM degradability, and other *in vivo* properties.^21, 22^

There are several types of 3D models in which cancer multicellular tumor spheroids (MCTS, which resemble OCAS) can be grown *in vitro*.^23^ During peritoneal metastasis, OvCa cells migrate into the peritoneal fluid and form OCAS.^3, 5^ These OCAS later attach to and invade surrounding tissues.^24^ For this reason, *in vitro* models of OvCa tissue coupled with patient-derived OCAS could improve OvCa drug discovery.^25^ Thus, they promise fewer false leads, and so better drug discovery. Moreover, the use of established cell lines does not capture disease heterogeneity across patients.^26^ While genetic mechanisms of OvCa have been studied,^27^ these findings alone have not been sufficient for explaining drug response.^28^ The use of primary cells isolated from patients would improve testing of drugs in a more personalized way.^29^ Ascites fluid from OvCa patients is drained from the peritoneal cavity from the patient to relieve the pain, this fluid is rich in single malignant cells and OCAS.^30, 31^ In this study, we evaluated the response of SKOV-3 OvCa MCTS and patient-derived OCAS to several drugs across different mechanisms of action in synthetic, tailorable hydrogels. Through this study we revealed a non-intuitive response of patient cells to these drugs, and we nominate collagen 3 as a particularly interesting ECM protein potentially driving resistance to platinum-based therapies in 3D. These hydrogels are easy to create and support the viability and survival of patient-derived OCAS cells, and we propose this as a useful system to pre-clinically screen drug candidates.

## Results

### OvCa MCTS were viable and proliferative in 3D poly (ethylene glycol)-maleimide (PEG-MAL) hydrogels

OvCa metastasizes to several sites within the peritoneal cavity.^4^ Here, we chose to focus on the omentum due to the availability of published data on this tissue of metastasis that we used to design our 3D hydrogel model. Since OvCa metastasizes to the omentum as aggregated cells,^31^ we hypothesized that a 3D hydrogel with integrin-binding peptides found in the omentum would support the viability and proliferation of OvCa MCTS (Figure 1). We previously used three methods for creating MCTS from established cell lines,^23^ and here we chose to use the microwell formation method, since this is more akin to how OvCa form OCAS (aggregated cells rather than clonal growth from a single cell) *in vivo* during metastasis.^31^ Additionally, our previous work demonstrated that this method is fast (MCTS form within one day), and resulted in the most uniform size distribution of MCTS.^23^ For this study we selected the widely used SKOV-3 OvCa cell line to form MCTS. We then chose to encapsulate these MCTS in 3D poly (ethylene glycol)-maleimide (PEG-MAL) hydrogels to capture a physiologically relevant stiffness with high water content, as well as tailored integrin-binding and matrix degradability. Since PEG hydrogels are not inherently degradable by cells, it was necessary to incorporate a cell-degradable crosslinking sequence during hydrogel synthesis to facilitate cell survival and spreading within the matrix. For this study, we selected a Pan-matrix metalloproteinase (Pan-MMP) degradable peptide crosslinker that can be degraded by many MMPs at the glycine-isoleucine bond.^32^ To optimize MCTS survival, we varied the amount of Pan-MMP degradable crosslinking within the hydrogel that contained RGD peptideintegrin-binding (Figure S1a). The total crosslinking was kept to a 1:1 molar ratio of maleimide-functional end groups to thiol-functional end groups by completing the crosslinking solution with linear PEG-dithiol (PDT) while the amount of Pan-MMP degradable crosslinking was varied. From our data we concluded that 12 molar % Pan-MMP degradable crosslinking supported MCTS survival and we used this condition for all subsequent experiments unless otherwise noted.

**Figure 1:**
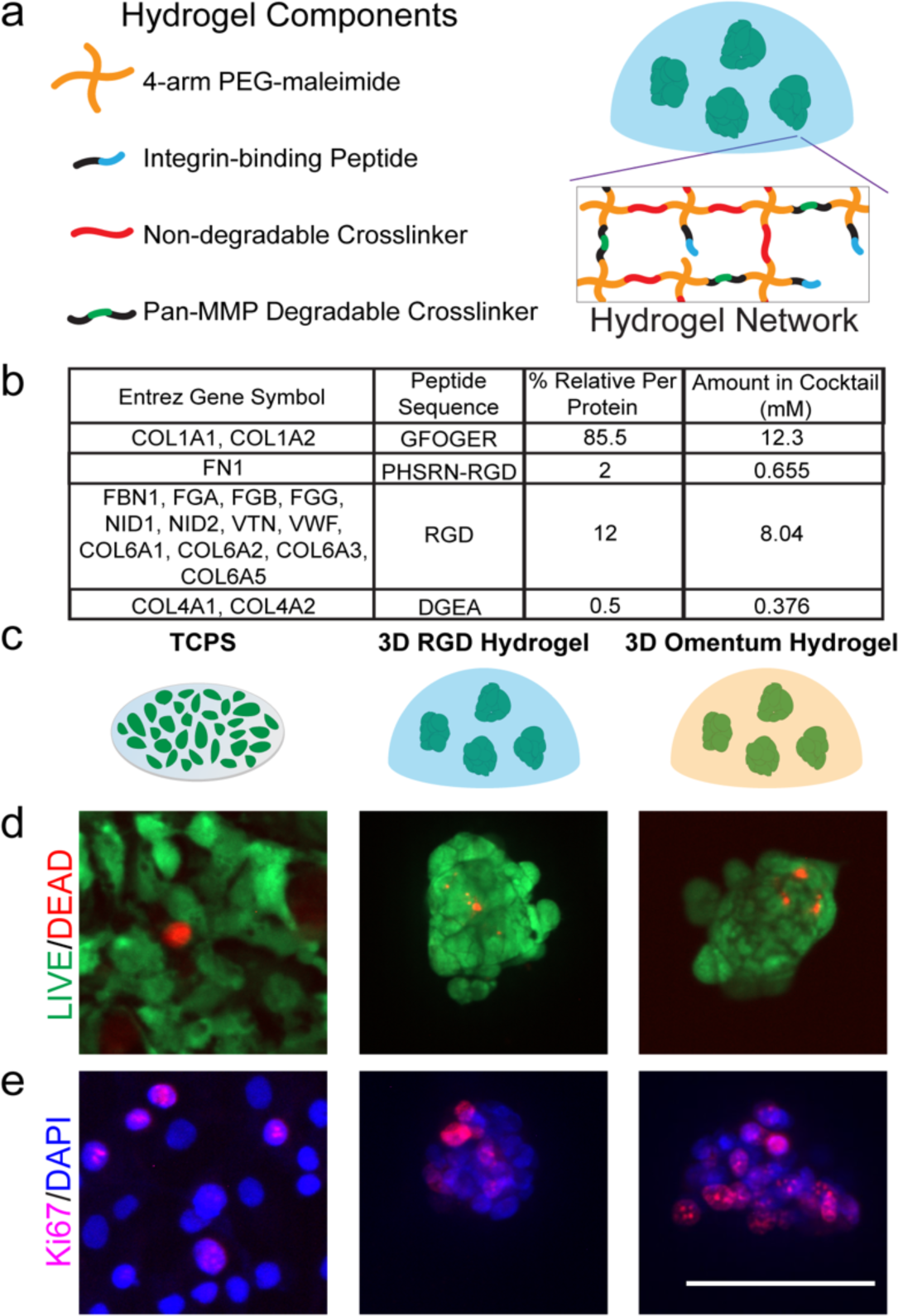
Three biomaterial platforms that were used to evaluate SKOV-3 drug response. a. Schematic of the components of a 3D PEG-MAL hydrogel with encapsulated SKOV-3 MCTS within a 3D hydrogel. b. Omentum-inspired integrin-binding peptide cocktail based on healthy omentum proteomic data from Naba et al.^33^ c. Schematic of cells grown on plastic or encapsulated in a 3D hydrogel model. d. LIVE(green)/DEAD(red) imaging of cells in each platform one day post-encapsulation. e. Ki67(pink)/DAPI(blue) staining of cells one day post-encapsulation to evaluate cell proliferation. Scale bar = 100 μm. Images are representative from N ≥ 2 independent experiments.

We tested encapsulating MCTS from one, two, or three wells of the 12-well plate containing microwells into six PEG-MAL hydrogels with RGD peptide integrin-binding and 25 molar % Pan-MMP degradable crosslinking with the remainder of the crosslinking with linear PDT to ensure that there were enough MCTS in each hydrogel for a detectable output signal for the drug screening assay. We found that the ratio of one well from the 12-well plate to six PEG-MAL hydrogels was ideal for a CellTiter-Glo output signal where we would be able to detect changes in the MCTS response with varying drug concentrations. Additionally, we found that the relative change in output signal from day one to day three was approximately the same for each ratio of wells from the microwells plate to six PEG-MAL hydrogels (Figure S1b). We used MCTS from one well of a 12-well microwells plate for six PEG-MAL hydrogels for the remainder of the study since it allowed us to test as many conditions as possible using one 12-well microwells plate and did not reduce the quality of the final assay output.

We measured the drug response of SKOV-3 MCTS in a 3D PEG-MAL hydrogel with RGD integrin-binding, but we hypothesized that adding tissue specificity could provide a better environment for MCTS survival and therefore the drug responses would be more representative of what would occur *in vivo*. We therefore increased the complexity of the 3D hydrogel model by adding additional integrin-binding peptides specific to the omentum, which is the site of many OvCa metastases.^4^ We aimed to create a 3D hydrogel with some tissue specificity without being overly complex that would limit future applications in drug screening or in a clinical setting. To find the most abundant integrin-binding proteins in the omentum, we mined published mass spectrometry data of healthy omentum tissue.^33^ The full integrin-binding peptide cocktail is described in Figure 1b. GFOGER, which is the binding sequence in collagen 1 and other fibrillar collagens^34^ comprises a majority of the hydrogel, but this cocktail contains RGD, as well as PHSRN-RGD and DGEA representative of other proteins described in Figure 1b. For simplicity, we kept the Pan-MMP degradable crosslinker used with the RGD only integrin-binding hydrogel to allow for MCTS to degrade the surrounding hydrogel.

Since this omentum-inspired hydrogel is new to the field, we first ensured that cells were viable and proliferative before drug screening, alongside two other platforms as internal controls (Figure 1c). As expected, SKOV-3 cells cultured on TCPS showed high viability and proliferation indicated by LIVE/DEAD and Ki67 staining, respectively (Figure 1d-e). Both of our PEG-MAL hydrogel models supported cell survival with very few dead cells inside the MCTS. By visual inspection of Ki67 stained images, there appeared to be a greater proportion of proliferative cells in the omentum hydrogel than on TCPS (Figure 1e). These results allowed for subsequent drug screening experiments in these platforms.

### SKOV-3 OvCa MCTS were resistant to Paclitaxel, and sensitized to other drugs in the 3D PEG-MAL hydrogels relative to TCPS

As a control, we first established drug responses of the SKOV-3 cells to five drugs, spanning different mechanisms of action, on TCPS (Figure 2, S2-3). We quantified the IC_50_, EC_50_, and GR_50_ ^35^ (Figure S2) for each cell line and drug combination using the online GR calculator.^36^ We calculated and reported these three parameters because they each provided different information about the potency of the drug on the cell line.^35^ In all cases, the GR_50_ was lower than the IC_50_ and EC_50_ metrics (Figure S2), because the GR_50_ is quantifying the reduction in *growth rate* and not overall inhibition of cell number. We found that SKOV-3 MCTS did not respond to Paclitaxel in the 3D hydrogels (EC_50_, Figure 2b, S4-5), but had an average EC_50_ of 12.1 μM on TCPS. We chose to report and compare the EC_50_ for each condition since it is the concentration that corresponds to achieve half of the maximal response and does not require at least 50% cell death as the IC_50_ does.^35^ This means that the drug responses across multiple conditions could still be compared if the cells did not grow exponentially over the course of the experiment or if greater the 50% cell death relative to the control was not achieved for which is required for calculating the GR_50_ and IC_50_, respectively. We found that it was not possible to calculate the GR_50_ in many of the 3D cases that we tested. Since Paclitaxel is a first-line therapy for OvCa patients, we were surprised to see a lack of response of SKOV-3 MCTS in our hydrogel models to this chemotherapy. However, this result has been previously reported in another 3D model by Loessner *et al*.^37^ This observation is important, because the drug response curve for SKOV-3 with Paclitaxel on TCPS is typical (Figure S3b), making it appear efficacious, but the curve looks strikingly dissimilar in a 3D culture model.

**Figure 2:**
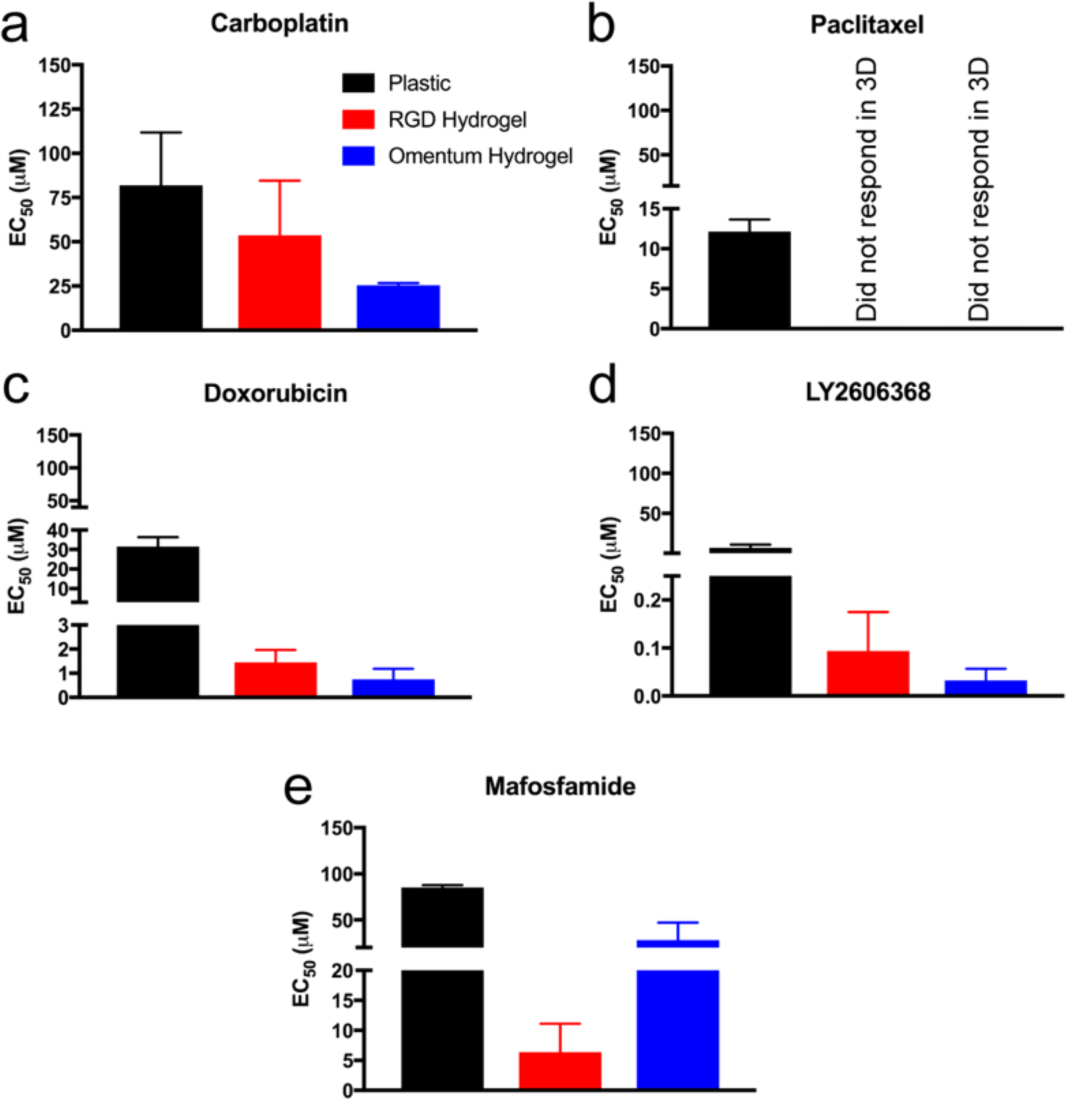
Quantified SKOV-3 drug response reported as EC_50_ on TCPS, or as MCTS encapsulated in RGD or omentum-inspired peptide-binding 3D PEG-MAL hydrogels. a. Carboplatin. b. Paclitaxel. c. Doxorubicin. d. LY2606368. e. Mafosfamide. Data are reported as the mean ± SEM. The EC_50_ values were calculated using the online GR calculator and from N ≥ 2 independent experiments with N = 3 technical replicates on TCPS and N ≥ 2 technical replicates in hydrogels.

Surprisingly, we found that the MCTS in the hydrogels were more sensitive to all the non-platinum drugs that we tested, relative to TCPS (EC_50_, Figure 2, S4-5). SKOV-3 MCTS were more sensitive (lower EC_50_) to Carboplatin and Doxorubicin (Figure 2a,c) in the omentum hydrogel than in the RGD hydrogel. On the other hand, MCTS were more sensitive to LY2606368 and Mafosfamide (Figure 2d-e) in the omentum hydrogel than on TCPS, although MCTS were less sensitive to Mafosfamide in the omentum hydrogel compared to the 3D RGD hydrogel;; *note* these compounds have not yet become clinically approved for OvCa treatment. An expanded future study with these models and other drugs, both clinically approved and in development, could possibly be used to make correlations between pre-clinical studies and clinical efficacy in patients.

### Patient-derived OCAS drug sensitivity reflected the real clinical response in omentum-mimicking hydrogels

Our goal was to evaluate how OCAS from patient samples responded to the drugs in the 3D omentum-mimicking hydrogel relative to TCPS. We collaborated with the team at UMass Medical School to obtain ascites fluid for screening OCAS. We then encapsulated OCAS into 3D omentum hydrogels, and compared their 3D drug responses to that of TCPS (Figure 3). The basic pathology reports for the patient samples have been listed in Figure 3a. Prior to drug screening, samples were assessed for viability via LIVE/DEAD staining (Figure 3b), which showed high viability across all samples in the 3D PEG-MAL omentum hydrogels. We observed that samples from Patient 1 were resistant to Mafosfamide, but sensitive to LY2606368 in the 3D hydrogel relative to TCPS (Figure 3c, S6), an inhibitor that is currently in Phase 2 clinical trials for OvCa.^38, 39^ However, upon receiving the next sample (Patient 1_2_ was the same as Patient 1 with six weeks between sample collection), resistance in 3D versus TCPS increased for these two drugs. Furthermore, this sample was resistant to all the drugs in 3D relative to TCPS, which demonstrated how resistant OvCa could be for current first line therapy, especially since this patient had received Paclitaxel treatment. Patient 2 received both Carboplatin and Paclitaxel (Figure 3a). Our 3D omentum hydrogel model demonstrated sensitivity to Carboplatin, Doxorubicin, LY2606368, and Mafosfamide relative to TCPS, and only slight resistance to Paclitaxel. If Patient 2, in reality, was not responding to these drugs in the clinic, and the response was not dependent on geometry, we expected to see similar responses on TCPS and in 3D. Interestingly, Doxorubicin, LY2606368, and Mafosfamide all appeared to be more effective in 3D versus 2D for the Patient 2 sample. The results from the samples with Patient 1 indicate that our hydrogel model may be a way to predict potential drug response as the disease progresses, which could be a way to help guide future clinical treatment decisions. However, more samples would be required to establish generalized observations of eventual prognostic value.

**Figure 3:**
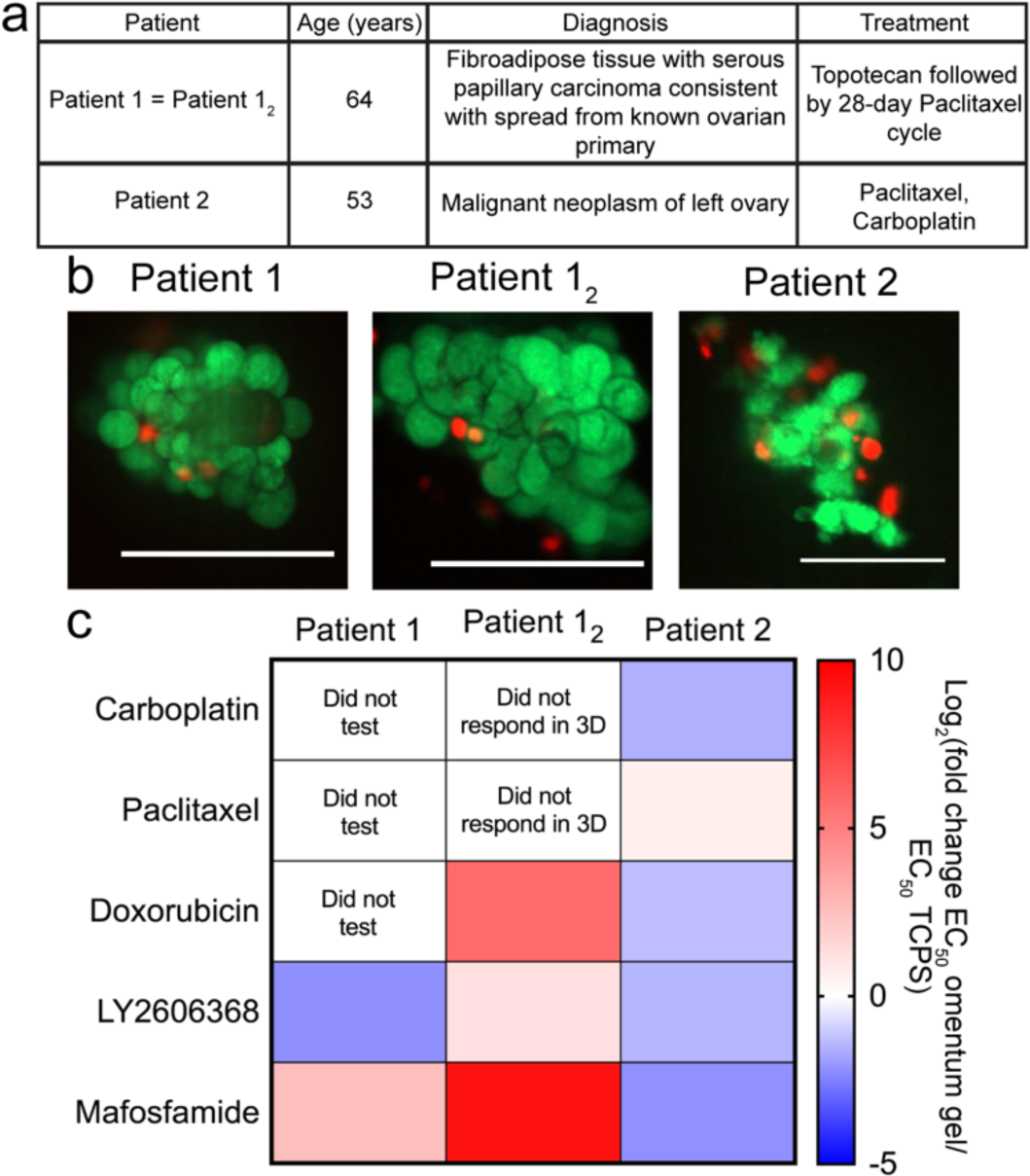
OvCa samples derived from patient ascites were encapsulated in 3D omentum-inspired PEG-MAL hydrogels for drug screening. a. Pathology report of patient samples. b. LIVE(green)/DEAD(red) imaging of OCAS in 3D omentum PEG-MAL hydrogels one day post-encapsulation. Scale bars = 100 μm. Images are representative from the sample. c. Patient sample drug responses displayed as log_2_ fold change in EC_50_ in the omentum hydrogel relative to the _50_ on TCPS. Red and blue indicate higher and lower, respectively, EC_50_ values in the 3D omentum hydrogel relative to the EC_50_ on TCPS.

### SKOV-3 cells expressed abundant collagen 3 protein in 3D hydrogels

To explore possible reasons for the differences in drug responses that we observed, we quantified the presence of a subset of ECM proteins via immunofluorescence staining (Figure 4). Fibronectin has been shown to be produced by mesothelial cells to promote OvCa metastasis,^40^ and we observed fibronectin production on TCPS and in 3D hydrogels by the OvCa cells themselves (Figure 4a). By visual inspection, in both 3D hydrogel models it appeared that the fibronectin was localized to within the MCTS rather than around the MCTS surface. Collagen 1 is a major component of the omentum microenvironment,^33^ and work by Januchowski *et al*. discovered fibrillar collagen genes, and, in particular, collagen 3A1 were upregulated in Topotecan- and Paclitaxel-resistant OvCa cells on TCPS.^41^ Therefore, we looked for both extracellular collagen 1 and collagen 3 (detected by collagen 1 and collagen 3A1 antibodies, respectively) in the environments tested here. We found that in the 3D hydrogels, but not on TCPS, SKOV-3 cells produced collagen 1 (Figure 4b). This suggests that perhaps the SKOV-3 MCTS were working to produce their own fibrillar collagen 1 for survival since GFOGER does not provide a fibrillar structure for the cells to most effectively bind. Furthermore, we observed some collagen 3A1 on TCPS on day 3, whereas it was highly expressed in both 3D conditions at both timepoints (Figure 4c). These results suggest that SKOV-3 cells may produce their own ECM proteins to increase their survival to prevent response to Paclitaxel exposure in 3D. Future work could examine a larger panel of ECM proteins to attempt at correlating the presence on specific proteins and OvCa drug resistance in 3D microenvironments.

**Figure 4:**
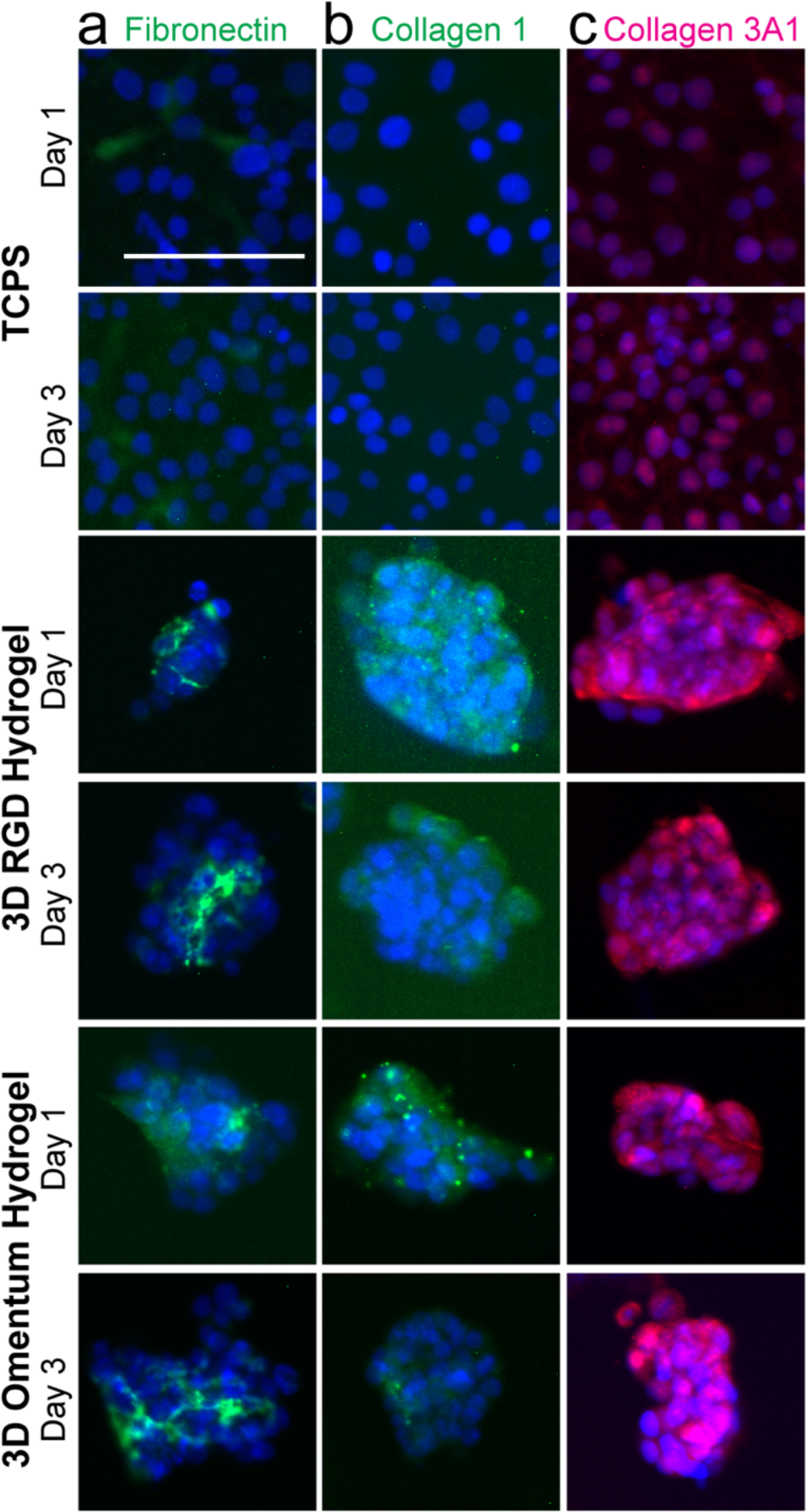
Immunofluorescence staining of SKOV-3 cells demonstrated that they produce their own ECM proteins over the time frame of a drug screening assay. a. Fibronectin(green)/DAPI(blue). Fibronectin can be seen on plastic as well as between cells in the MCTS. b. Collagen 1(green)/DAPI(blue). Collagen 1 production was not observed on plastic, but was visible around the spheroids. c. Collagen3A1 (pink)/DAPI (blue). Some collagen 3A1 may be present on plastic, but is clearly visible surrounding the spheroids in both hydrogels. Scale bar = 100 μm. Images are representative from N ≥ 2 independent experiments.

## Discussion

There is clear room for improvement in OvCa treatment. The availability of ascites from OvCa patients provides a valuable opportunity for developing an improved OvCa drug testing pipeline that combines patient cells and engineered environments. Others have recognized the need for developing more clinically relevant OvCa drug screening platforms.^37, 42^ Similar to our RGD hydrogel, Loessner *et al*. evaluated the response of SKOV-3 MCTS to Paclitaxel in a 3D PEG-based hydrogel with RGD binding sites and MMP-degradable sites.^37^ The MCTS generation approach was different than ours where their platform allowed for the formation of the MCTS from single cells within the PEG hydrogel over 12 days rather than by the aggregation and encapsulation method that we used.^37^ Even with a different 3D hydrogel and MCTS formation method, they also observed resistance to Paclitaxel in 3D.^37^ However, they did not test the response to other drugs in their study, which limits what we can compare between studies. Liu *et al*., with 3D collagen gels, observed resistance of OV-NC and OV-206 OvCa cell lines seeded as single cells in 3D to Paclitaxel, Carboplatin, and 5-fluorouracil relative to TCPS.^43^ Furthermore, the use of this collagen gel demonstrated that the ability of OvCa to interact with the collagen matrix increased the invasive capacity of the cells.^43^ From this observation and our collagen staining results, it appears that collagen 1 is necessary for OvCa disease progression and/or drug resistance in 3D, and OvCa cells will produce this protein themselves if it is not provided within the biomaterial environment. To model the omentum for understanding OvCa metastasis, a multicellular platform containing mesothelial cells, fibroblasts, and ECM was developed.^44^ The application of this platform identified that mesothelial cells from the omentum caused inhibition and fibroblasts and ECM promoted attachment and invasion of OvCa cells.^44^ In the context of OvCa drug screening and the omentum microenvironment, we see our study as a valuable contribution to OvCa drug response research because we compare OCAS derived from ascites to cell line MCTS help establish links between pre-clinical drug screening models and clinical response.

We suggest that our synthetic hydrogel model is a good tool for screening patient OCAS drug response. By controlling the polymer concentration of the 3D hydrogels we were able to test the samples at a stiffness that is physiologically relevant to the omentum.^21, 45^ Furthermore, the 3D PEG-MAL hydrogels are easy to make because all of the peptides and polymer precursors can be made in advance and simply require dissolution for sample encapsulation. Thus, whenever patient samples are available, one can process the sample and complete the encapsulation within a few hours. If multiple treatment options are being considered for a patient, then this could be a simple, but valuable tool in guiding the selection of the next treatment steps. For example, LY2606368 is currently in Phase II clinical trials^38, 39^ and could be an option for patients in the future. Testing this compound in 3D hydrogels demonstrated it was more promising as a patient treatment than screening only on TCPS by comparing the reported EC_50_ values (Figure 3c). Furthermore, patient sample 1_2_ was received about 6 weeks after patient sample 1, and we observed an increase in resistance to the drugs that we tested. This implies that as disease progresses, the patient’s ability to respond to treatment decreases. To our knowledge, Mafosfamide has never been tested in clinical trials for OvCa, but has shown promise *in vitro* on TCPS.^46, 47^ The results from our 3D hydrogel approach suggest that this drug could be beneficial for patients, and this prediction of efficacy was likely missed with earlier studies that relied solely on TCPS (Figure 3c, S6). If the additional integrin-binding peptides of the omentum hydrogel more faithfully capture the *in vivo* microenvironment, then this could potentially help explain how some compounds have been able to successfully become clinical OvCa treatments. However, it should be noted that the added complexity of additional integrin-binding peptides in the omentum hydrogel was not necessary for observing the SKOV-3 resistant to Paclitaxel in 3D. From our experiments, however, it is not clear if the 3D MCTS structure or if the 3D hydrogel environment is driving this lack of response across the concentrations of Paclitaxel what we tested. Overall, we suggest that our approach could *and should* be extrapolated to more possible OvCa drugs either originally overlooked or currently under consideration/development.

In our previously published work, we compared differences in gene expression of MCTS spheroids created with a variety of different methods (i.e. clonal growth in polyNIPAAM vs. aggregation in microwells).^23^ In this first generation approach, we created MCTS by clonal growth from SKOV-3 cells and compared their response and the response of OCAS from patients to a small panel of drugs in a non-degradable RGD integrin-binding 3D synthetic PEG hydrogel.^23^ The results suggested further exploration of OvCa in a 3D model may be useful for understanding drug response in patient samples. Here we used microwells, an aggregation formation method, to create MCTS, since OCAS form by aggregation in patients.^31^ By simply encapsulating the cells and OCAS from ascites we preserved the cell-cell contacts and any associated ECM directly from the patient. Because we were limited in sample sizes, we are unsure of how much ECM comes with the OCAS, but we observed that the SKOV-3 cell line secreted fibronectin, collagen 3, and collagen 1 within one day of encapsulation (Figure 4). Since these ECM proteins were not seen on TCPS for all of these cases, we suspect that ECM production facilitated survival in 3D by adding new integrin binding sites in addition to those we provided via peptides. Our results aid in identifying collagen 3 as an important contributor to OvCa drug resistance since it was identified to be co-expressed with the ECM protein lumican in a Topotecan-resistant OvCa cell line on 2D by another research group.^48^ Perhaps this suggests future work could potentially explore how these ECM proteins directly impact OvCa response to drug.

It has been well-established that cells grow at different rates in 2D and 3D cultures.^12, 49^ Since the ability of cells to respond to chemotherapy is proliferation-dependent, response is expected to vary depending on growth conditions. To account for differences in growth rates that will impact traditional drug screening metrics, the GR_50_ was established.^36^ One key condition that must be met when applying this analysis is that the cells are growing exponentially over the course of the assay.^36, 50^ We have found that this key condition is not always met, especially in biomaterial platforms, and have discussed this in further detail.^35^ In this study, we were not able to calculate the GR_50_ for all of the conditions that we tested due to growth limitations. Therefore, the EC_50_ is what we were able to report here for all of the conditions that we tested. Due to the differences in growth rates of cells in 2D and 3D, our 3D model helps aid in understanding growth of the MCTS in 3D *in vitro* before moving to *in vivo* studies. We tested drugs used clinically to treat patients, a drug that is in clinical trials, and Mafosfamide, which has shown promise *in vitro* on TCPS,^46, 47^ but to our knowledge, has never been studied in clinical trials. As more becomes known about how the microenvironment impacts drug response, it may be possible to evaluate additional quantitative phenotypes that would aid in comparing drug responses across different geometries. Others have created models with other cell types including mesothelial cells and fibroblasts,^44, 51, 52^ but here our system was focused on the parameters of integrin binding sites, stiffness, and 3D geometry without the contribution of other cells present within the omentum microenvironment. Since ascites is rich in cytokines and growth factors,^30^ perhaps future studies with patient derived OCAS could incorporate ascites-derived cytokines as an additional feature for more faithfully modeling the omentum microenvironment for OvCa drug screening. By evaluating the presence of a subset of ECM proteins in addition to the EC_50_ for each condition, we have enabled an approach to couple the microenvironment composition and drug response, which could be applied in other drug screening studies with biomaterials. This could help establish large drug response and ECM composition data sets, which could then possibly be used to determine correlations between ECM composition and drug response that may aid in making better clinical predictions for the success of a drug candidate.

OvCa is a challenging disease to treat and development of second line therapies is critical, since recurrence occurs in a majority of patients. We propose that this can be achieved by using a 3D PEG-MAL hydrogel with encapsulated MCTS to aid in making better pre-clinical predictions for new therapeutic compounds. Our results demonstrated that screening for therapies on TCPS and in 3D hydrogels did not yield the same drug responses. This observation may be dependent on the ECM composition, and we concluded that TCPS may not be sufficient for evaluation of OvCa drug response. Most patients have the presence of ascites, which is removed as part of treatment and contains many OCAS. The integration of OCAS in an omentum-inspired 3D hydrogel model to study the drug response their drug response yielded results also different than on TCPS. We discovered that our 3D hydrogel model demonstrated drug response results that we could not have captured on TCPS alone, and recommend that future OvCa drug screens employ our materials to aid in understanding OvCa response *in vitro*.

## Methods

### Cell culture

All cells were maintained at 37°C and 5% CO_2_, and all cell culture supplies were purchased from Thermo Fisher Scientific (Waltham, MA) unless otherwise noted. The SKOV-3 cell line was purchased from the American Type Culture Collection (ATCC, Manassas, VA). SKOV-3 cells were grown in Roswell Park Memorial Institute (RPMI) medium supplemented with 10% fetal bovine serum (FBS) and 1% penicillin/streptomycin (pen/strep).

### Primary ovarian cancer ascites culture

Ascites samples were received from patients undergoing paracentesis at UMass Medical School (Worcester, MA), transported to UMass Amherst the same day as the procedure, and used for experiments immediately upon receipt. Samples were deidentified and were IRB exempt. Pathology reports are provided in Figure 3a. Single cells and OCAS were recovered from patient samples and treated as described previously.^23^ Samples containing both single cells and OCAS were grown on 2D TCPS or encapsulated directly into 3D PEG-MAL hydrogels. All experiments with primary OvCa samples were conducted as one biological replicate with RPMI + 10% FBS + 1% pen/strep.

### Microwell MCTS

Square pyramidal microwells (400 μm side-wall dimension) were fabricated as described previously.^53^ Briefly, master molds containing square-pyramidal pits were generated by anisotropic etching of 100 crystalline silicon in potassium hydroxide (KOH). Microwells were generated from poly(dimethylsiloxane) (PDMS) using a two-stage replica molding process of the master mold as described previously.^53^ Microwells were arranged in a square array with no space between adjacent wells and placed in 12-well plates. For cell seeding, microwell surfaces were UV sterilized for 30 minutes, pretreated with 5% Pluronic F-127 (Sigma-Aldrich, St. Louis, MO) for 5 minutes at 3,500 RPM (2,465 x g, Eppendorf 5810R v3.3 centrifuge with A-4-62 rotor, Eppendorf, Hamburg, Germany) with an additional 30-minute incubation on the bench, and then washed twice with sterile water. Cells were seeded 1.00 × 10^5^ cells/well (approximately 26,000 cells/cm^2^). After 24 hours, MCTS were collected by shaking the plate gently to dislodge most of them, and gently aspirating medium and MCTS. MCTS solution was spun down at 400 RPM (Thermo Scientific Sorvall ST16R centrifuge) for 5 minutes. Medium was removed and MCTS were encapsulated in 3D hydrogels and RPMI + 5% FBS + 1% pen/strep was added to each well after the hydrogels had polymerized.

### 3D PEG-MAL hydrogel platform

MCTS or OCAS were encapsulated into 3D PEG hydrogels as described previously^23^ with minor modifications. The 3D RGD hydrogel was prepared with a 20 kDa 4-arm PEG-maleimide (PEG-MAL, Jenkem Technology, Plano, TX) at 10 wt % solution (measured Young’s modulus at this condition^21^ corresponds with initial average measured elastic modulus of omental tissue^45^) with 2 mM of cell adhesion peptide GSPCRGDG (RGD, Genscript, Piscataway, NJ) and crosslinked with a 90 mM combination of the 1 kDa linear PEG-dithiol (JenKem, 79.2 mM) and cell-degradable cross-linker, Pan-matrix metalloproteinase (Pan-MMP, GCRDGPQGIWGQDRCG, Genscript, 10.8 mM)^32^ in sterile 2 mM triethanolamine (pH 7.4).The hydrogels were synthesized by pipetting 1 μL of PDT and Pan-MMP solution onto the well surface of a 48-well TCPS plate, followed by pipetting 9 μL of the MCTS/OCAS-containing PEG−MAL solution. The 3D omentum hydrogel was prepared with 2 mM of the omentum integrin-binding peptide cocktail. The omentum cocktail is fully described in Figure 1b and consists of (GRGDSPCG, Genscript), GFOGER (CGP(GPP)5GFOGER(GPP)_5_, Genscript), PSHRN-RGD (CGPHSRNGGGGGGRGDS, same methods as described previously^21^), DGEA (GCGDGEA, same methods as described previously^21^). The MCTS from microwells were transferred to either a 3D RGD hydrogel or 3D omentum hydrogel at a ratio of one well of a 12-well microwells plate to six 10 μL PEG-MAL hydrogels. The PEG-MAL solution containing MCTS was transferred with cut pipette tips to minimize shear stress.

### Drug Screening

Single SKOV-3 cells were seeded in RPMI with 5% FBS and 1% pen/strep at 6,250 cell/cm^2^ on TCPS 96-well plates. SKOV-3 MCTS were recovered and encapsulated in 3D hydrogels with the same medium as TCPS. Samples collected from ascites were seeded in RPMI with 10% FBS and 1% pen/strep at the same density on TCPS. Drugs were added after 24 hours in the same type of medium that the cells were seeded in and the cells were incubated with drugs for 48 hours. Carboplatin (Tocris Bioscience, United Kingdom) was added in 10-fold serial dilutions at concentrations of 2×10^−5^ to 2×10^2^ μM. Paclitaxel (MP Biomedicals, Santa Ana, CA), Mafosfamide (Niomech, Germany), and Doxorubicin (LC Laboratories, Woburn, MA) were added in 10-fold serial dilutions at concentrations of 1×10^−5^ to 1×10^2^ μM. LY2606368 (AbovChem LLC, San Diego, CA) was added in 10-fold serial dilutions at concentrations of 3 × 10^−5^ to 3 × 10^1^ μM. Dimethyl sulfoxide (DMSO, Sigma-Aldrich) was used as a vehicle control for all drugs except carboplatin was dissolved in water.^54^ Cell viability was assayed at 24 hours post-cell seeding in a control plate and after 48 hours of drug incubation using the CellTiter-Glo luminescent viability assay (Promega, Madison, WI). After five (TCPS) or 20 (hydrogels) minutes after adding the CellTiter-Glo, luminescence values were measured in a BioTek Synergy H1 plate reader (BioTek, Winooski, VT). The drug response metrics were calculated using the online GR calculator^36^ found at http://www.grcalculator.org/grcalculator/ for each cell line and drug combination. However, the EC_50_ values for LY2606368 drug responses of Patient 1 and the second biological replicate of SKOV-3 MCTS in the omentum-inspired hydrogel were calculated using GraphPad Prism v7.0c (GraphPad Software, Inc., LaJolla, CA) due to limitations of the GR calculator with the drug response curves in these cases. Conditions that were reported as “did not respond in 3D” were determined by visual inspection of the drug response curves. The SKOV-3 data reported are from N ≥ 2 independent experiments with N ≥ 2 technical replicates in each experiment.

### Immunofluorescence (IF) staining

All antibodies and corresponding dilutions used for IF staining are listed in Table S1. Single cells seeded on TCPS or MCTS encapsulated in 3D hydrogels for 1 or 3 days were assessed for proliferation via Ki67 immunofluorescence. For the Ki67 staining, samples were rinsed with PBS, fixed with 10% formalin for 15 minutes, permeabilized with tris-buffered saline (TBS) containing 0.5% Triton X-100 (Promega), and blocked with AbDil (2 wt % bovine serum albumin (BSA, Thermo) in TBS with 0.1% Triton X-100, TBS-T) for 20 minutes. Samples were incubated for 1 (TCPS) or 2 (hydrogels) hours at room temperature with the primary antibody, rinsed three times with TBS-T, and incubated with a secondary antibody for 1 (TCPS) or 2 (hydrogels) hours. To stain for ECM proteins, the cells were incubated with the antibodies listed in Table S1 for the incubation times described above. Cell nuclei were labeled with DAPI (Thermo) at 1:1,000 in phosphate buffered saline (PBS) for five minutes and samples were rinsed three times with PBS. Fluorescence imaging was obtained on a Zeiss Spinning Disc Cell Observer SD (Carl Zeiss AG, Oberkochen, Germany). The brightness and contrast were adjusted post-image collection and SKOV-3 images are representative from N ≥ 2 independent experiments.

### LIVE/DEAD cell viability staining

Cells seeded on TCPS or MCTS encapsulated in 3D hydrogels for 1 and 3 days, were assessed for viability with LIVE/DEAD staining (L3224, Thermo) in accordance with the manufacturer’s instructions. Imaging was obtained on a Zeiss Spinning Disc Cell Observer SD (Zeiss). The brightness and contrast were adjusted post-image collection and SKOV-3 images are representative from N ≥ 2 independent experiments.

## Supporting information

All Supplemental Materials

## Acknowledgements

The authors are grateful to Dr. Kathleen Arcaro and Dr. Jungwoo Lee for use of equipment. Also, we would like to thank Hong Bing (Amy) Chen, Chien-I (Mike) Chang, and Cristian Fraioli at the Cancer Center Tissue and Tumor Bank at UMass Medical School for primary ovarian cancer ascites. We would like to thank Sualyneth Galarza for helpful discussions about designing the omentum peptide cocktail. This work was funded by a grant from the NIH (1DP2CA186573-01) and a CAREER grant from the NSF (1454806) to SRP. EAB was partially supported by a fellowship from the National Institutes of Health as part of the University of Massachusetts Chemistry-Biology Interface Training Program (National Research Service Award T32 GM008515). SRP is a Pew Biomedical Scholar supported by the Pew Charitable Trusts. SRP and MFG were partially supported by a Barry and Afsaneh Siadat Career Development Award.

## Author contributions

EAB and MFG performed all of the cell experiments. DCC and KRS designed and created the microwells. SRP supervised the study. EAB, MFG, and SRP wrote the manuscript.

