## Supplementary material for "An Omentum-inspired 3D PEG Hydrogel for Identifying ECM-drivers of Drug Resistant Ovarian Cancer": All Supplemental Materials

**Table S1: Antibodies used for immunofluorescence staining.**

| Target Protein | Primary Antibody | Primary Antibody Dilution in AbDil | Secondary Antibody | Secondary Antibody Dilution in AbDil |
|---|---|---|---|---|
| Ki67 | ab16667, Abcam, UK | 1:200 | Alexa Fluor 647 goat anti-rabbit, Molecular Probes (Eugene, OR) | 1:500 |
| Fibronectin | 563100, BD Biosciences, San Jose, CA | 1:200 | Alexa Fluor 488 was already conjugated to the primary antibody, Molecular Probes | N/A |
| Collagen 1 | ab6308, Abcam | 1:200 | Alexa Fluor 555 goat anti-mouse, Molecular Probes | 1:500 |
| Collagen 3A1 | A3975, ABclonal Technology, Woburn, MA | 1:200 | Alexa Fluor 647 goat anti-rabbit, Molecular Probes | 1:500 |

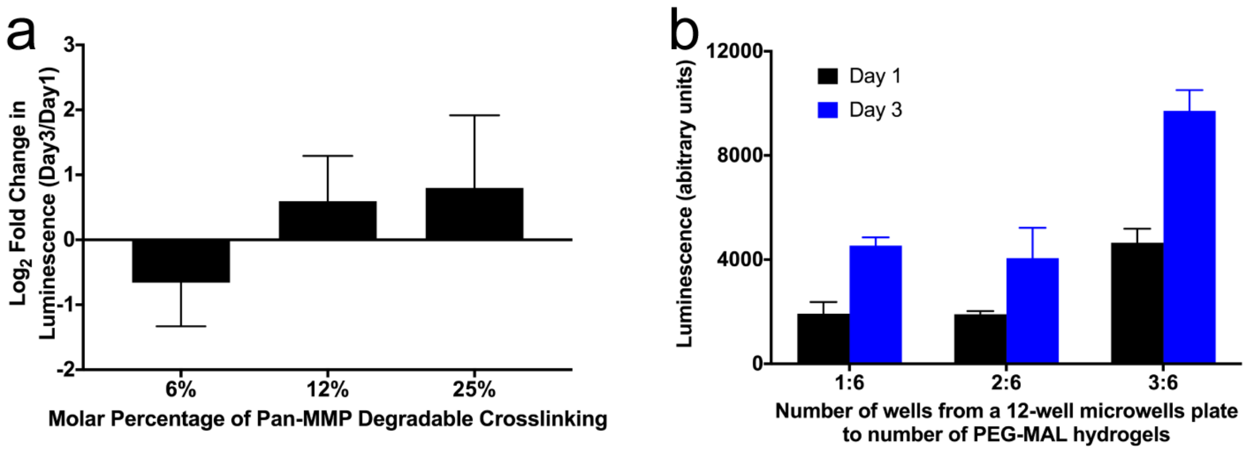


**Figure S1: Optimization of RGD integrin-binding 3D PEG-MAL hydrogels for SKOV-3 MCTS drug screening experiments** a. The effect of changing the molar % of Pan-MMP degradable crosslinking on MCTS survival reported as $\log_2$ fold change in luminescence measured from the CellTiter-Glo assay at day three relative to the average luminescence at day one. b. Luminesce tested during encapsulation of MCTS from one, two, or three wells of the 12-well plate containing microwells into six PEG-MAL hydrogels at day 1 and 3. Data are reported as mean ± SEM. N = 3 technical replicates.

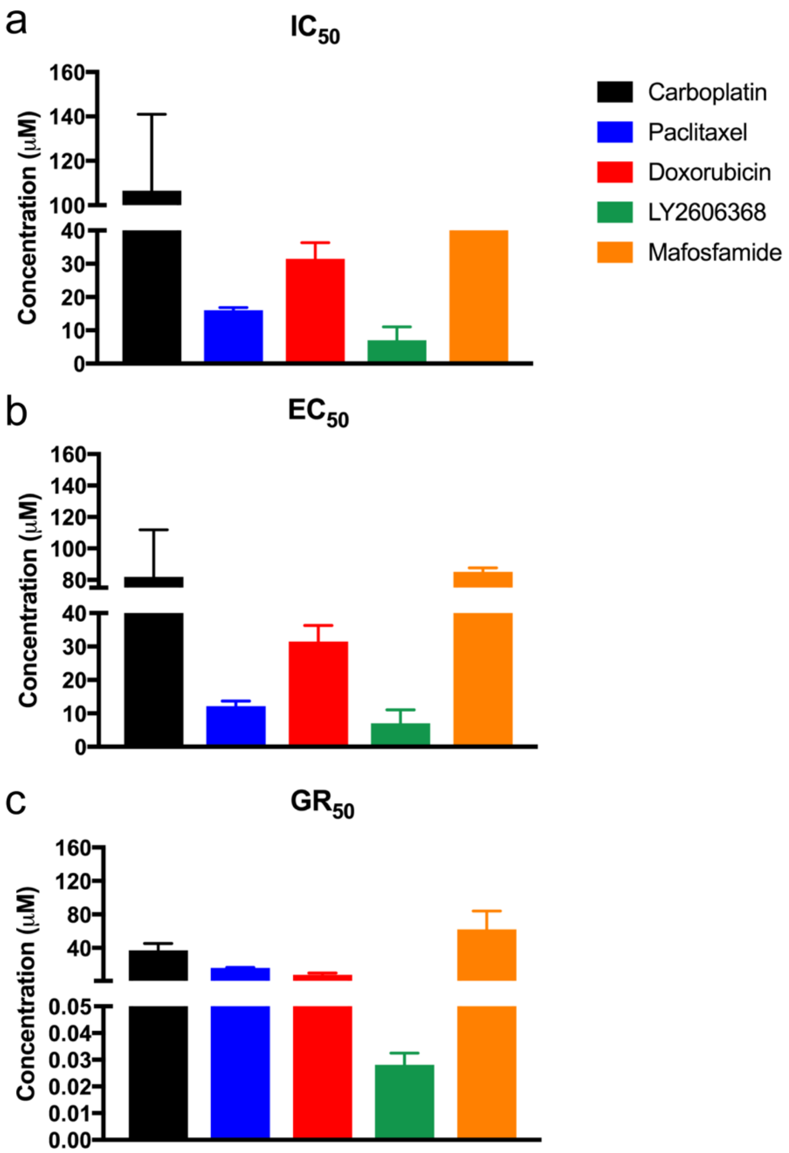


**Figure S2: Calculated drug screening metrics for SKOV-3 cells as a monolayer on TCPS.** a. $IC_{50}$. b. $EC_{50}$. c. $GR_{50}$. Data are reported as the mean ± SEM. N ≥ 2 two independent experiments with N = 3 technical replicates per experiment. Only one $IC_{50}$ could be calculated for Mafosfamide from the drug response curves.

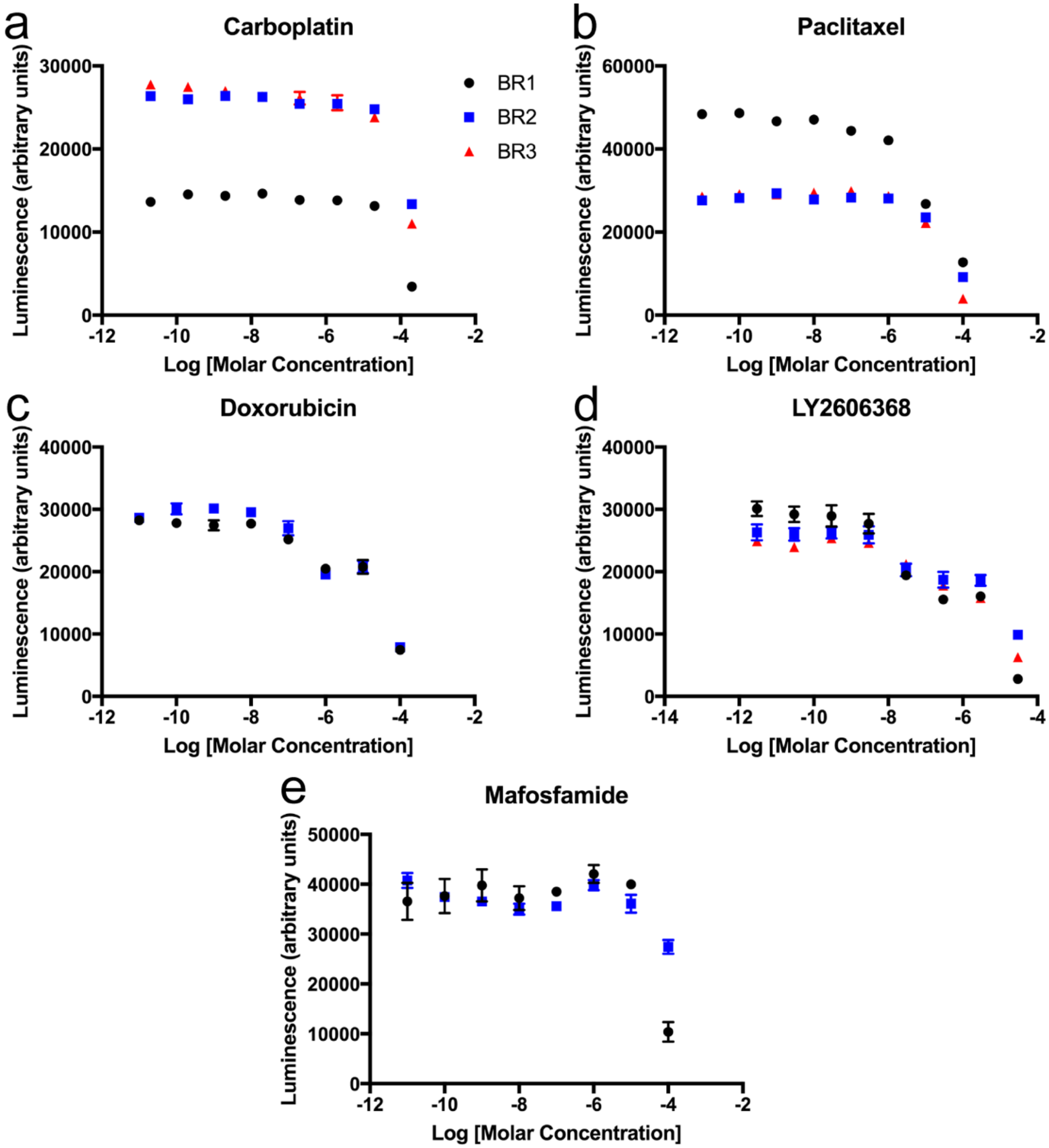


**Figure S3: Drug response curves at 48-hours post-dosing for SKOV-3 cells as a monolayer on TCPS measured with CellTiter-Glo.** a. Carboplatin. b. Paclitaxel. c. Doxorubicin. d. LY2606368. e. Mafosfamide. Data are reported as the mean ± SEM. N ≥ 2 two independent experiments (BR) with N = 3 technical replicates per experiment.

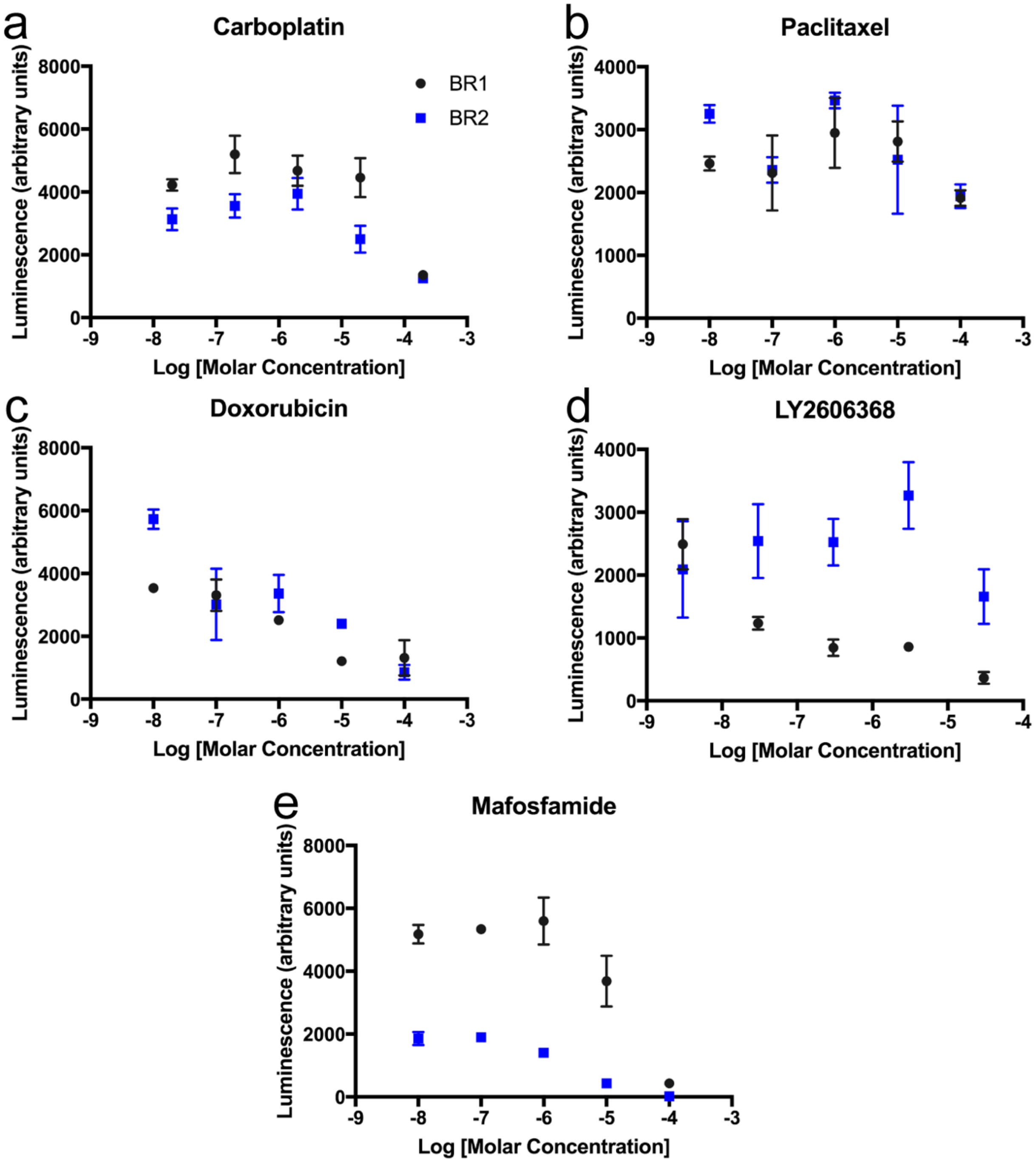


**Figure S4: Drug response curves at 48-hours post-dosing for SKOV-3 MCTS in 3D RGD integrin-binding PEG-MAL hydrogels measured with CellTiter-Glo.** a. Carboplatin. b. Paclitaxel. c. Doxorubicin. d. LY2606368. e. Mafosfamide. Data are reported as the mean ± SEM. N = 2 two independent experiments (BR) with N ≥ 2 technical replicates per experiment.

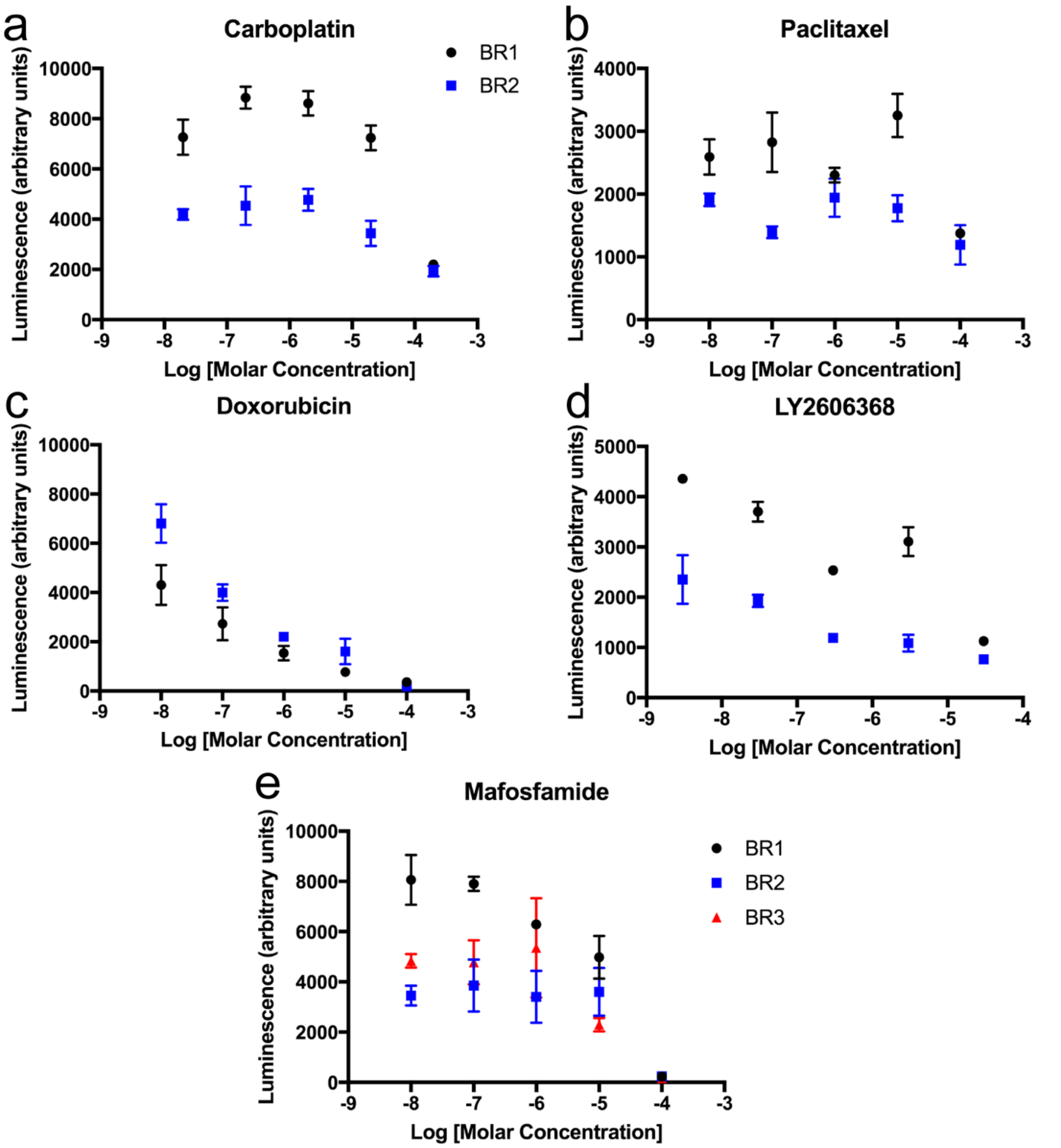


**Figure S5: Drug response curves at 48-hours post-dosing for SKOV-3 MCTS in 3D omentum-inspired integrin-binding PEG-MAL hydrogels measured with CellTiter-Glo.** a. Carboplatin. b. Paclitaxel. c. Doxorubicin. d. LY2606368. e. Mafosfamide. Data are reported as the mean ± SEM. N ≥ 2 two independent experiments with N ≥ 2 technical replicates per experiment.

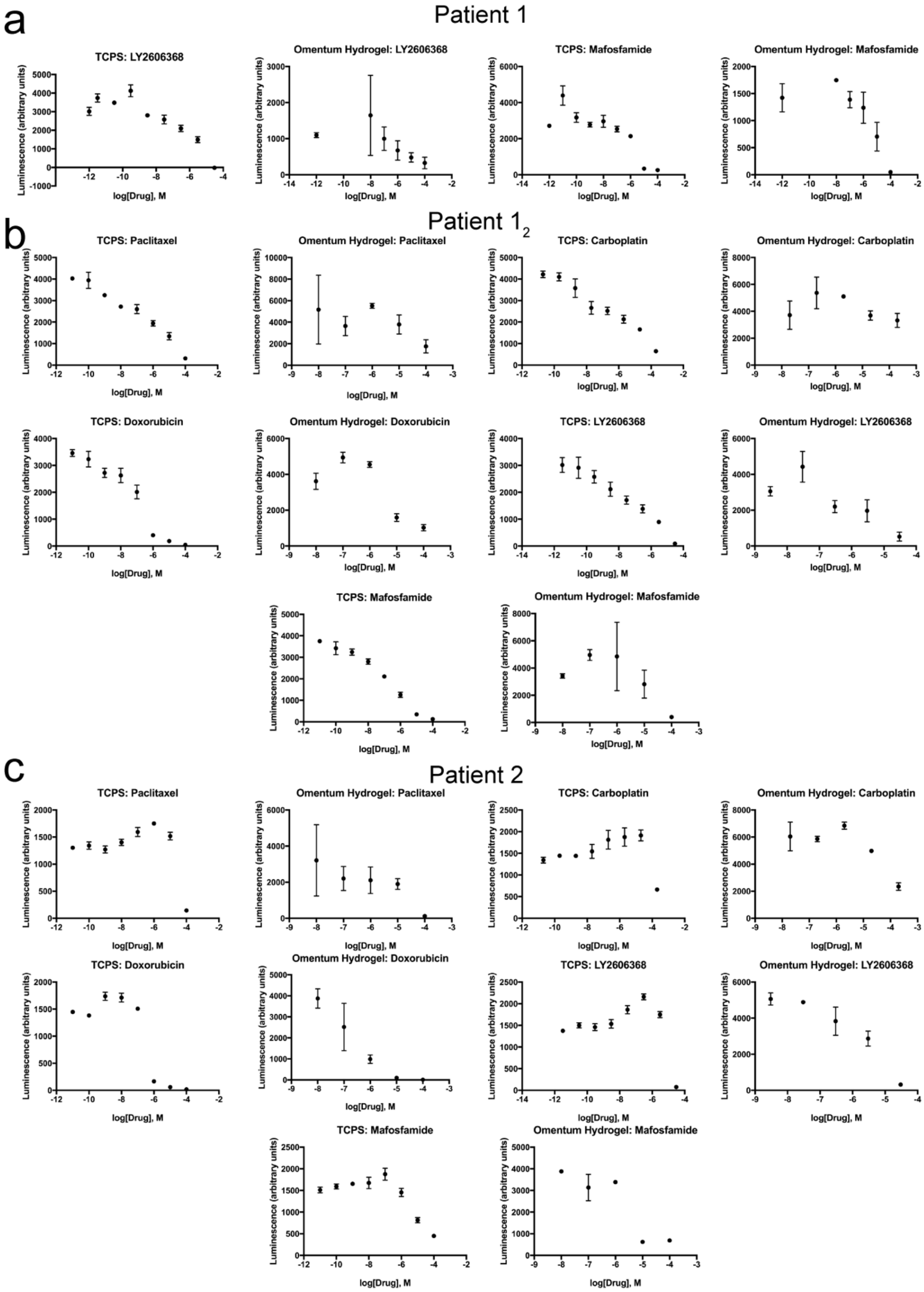


**Figure S6: Drug response curves at 48-hours post-dosing for ascites-derived patient samples on TCPS or in 3D omentum-inspired PEG-MAL hydrogels.** a. Patient 1. b. Patient 1$_2$. c. Patient 2. Data are reported as the mean ± SEM. N = 3 technical replicates on TCPS and N ≥ 2 technical replicates in 3D hydrogels.
